# MQuad enables clonal substructure discovery using single cell mitochondrial variants

**DOI:** 10.1101/2021.03.27.437331

**Authors:** Aaron Wing Cheung Kwok, Chen Qiao, Rongting Huang, Mai-Har Sham, Joshua W. K. Ho, Yuanhua Huang

## Abstract

Mitochondrial mutations are increasingly recognised as informative endogenous genetic markers that can be used to reconstruct cellular clonal structure using single-cell RNA or DNA sequencing data. However, there is a lack of effective computational methods to identify informative mtDNA variants in noisy and sparse single-cell sequencing data. Here we present an open source computational tool MQuad that accurately calls clonally informative mtDNA variants in a population of single cells, and an analysis suite for complete clonality inference, based on single cell RNA or DNA sequencing data. Through a variety of simulated and experimental single cell sequencing data, we showed that MQuad can identify mitochondrial variants with both high sensitivity and specificity, outperforming existing methods by a large extent. Furthermore, we demonstrated its wide applicability in different single cell sequencing protocols, particularly in complementing single-nucleotide and copy-number variations to extract finer clonal resolution. MQuad is a Python package available via https://github.com/single-cell-genetics/MQuad.

## Introduction

Identification of clonal relationships in a population of single cells is a major grand challenge of single cell data science^1^. Such information is essential in recovering cell lineages, which can have broad applications in developmental and stem cell biology, and cancer biology. In particular, deciphering intra-tumor genetic heterogeneity and clonal mutations are cricitial for revealing their evolutionary dynamics and drug resistance of cancers^2^. Fortunately, recent advances in single-cell sequencing bring promises to the identification of subclonal structure in tumors^3,4^ and characterization of phenotypic impacts from single nucleotide variations (SNVs) and copy number variations (CNVs)^5,6^. However, SNV-based lineage reconstruction remains a grand challenge^1^ partly due to the relatively small number of somatic variants in the large nuclear genome^7^ and a high dropout rate^8^, especially for assays with low efficiency and limited coverage, e.g. Smart-seq2 for transcriptome^9^. CNVs inferred based on single cell transcriptomic data have been widely used^10,11^, but subclonal structures inferred from read-coverage-inferred CNVs might not be as dynamic as subclones shown by the propagation of true point mutations. It has been shown recently that mitochondrial heteroplasmy serves as an excellent alternative to nuclear SNVs for studying lineages^12,13^ due to the mitochondria’s large number of copies and higher mutation rate (>10 fold of nuclear genome^14^). Thanks to its cost efficiency and high accessibility, mtDNA variations have since attracted great attention, with further development of novel droplet-based single-cell sequencing protocols to enrich mitochondrial sequences in a highly multiplexed manner, such as mtscATAC-seq^12^.

However, it is highly challenging to differentiate between clone-discriminative mtDNA mutations and non-inherited mutations that are totally unrelated to the lineage structure. Very few computational methods are available for analyzing mitochondrial SNVs across different sequencing assays, especially for conventional single-cell RNA sequencing (scRNA-seq) data. Nuclear SNV callers such as Monovar ^15^ and Conbase ^16^, assume a diploid context which is violated in the mitochondrial genome. MtDNA specific methods, EMBLEM^17^ and mgatk^12^ were designed primarily for single-cell assay for transposase-accessible chromatin with sequencing (scATAC-seq) and the SNV quality detected in other data types such as scRNA-seq is found to be not as robust, as stated by the developers of mgatk as one of its limitations^12^. Combining with the long standing problem of sequencing errors, uninformative and noisy mtDNA SNVs greatly confound clonal inference and biological interpretation.

To address these limitations, we developed a new computational method called MQuad (Mixture Modelling of Mitochondrial Mutations, M^4^) that effectively identifies informative mtDNA variants in single-cell sequencing data for clonality inference. Importantly, MQuad can be used in combination with two other recently developed tools to form a novel integrated clonality discovery pipeline, cellSNP-MQuad-VireoSNP, which provides a complete analysis suite from single cell mtDNA genotyping to clonal reconstruction. We demonstrate its usage on various single-cell sequencing data sets to identify clones based on mtDNA mutations. More importantly, our analysis reveals that mtDNA mutations detected by MQuad can be used in complement with nuclear SNVs and CNVs to achieve finer clonal resolution.

## Results

### MQuad is a robust statistical approach to identify informative mtDNA variants

MQuad is a computational method for detecting clone discriminative mitochondrial variants. It is tailored to work seamlessly with cellsnp-lite^18^ and vireoSNP^19^ to create an automated end-to-end pipeline for single cell clonal discovery using mitochondrial variants (Fig. 1a). Briefly, MQuad fits the alternative and reference allele counts of each variant to a binomial distribution with either one shared parameter (i.e., the expected alternative allele frequency) across all cells under the null hypothesis H_0_ or two different parameters in the cell population as the alternative hypothesis H_1_ (i.e., two-component mixture; Methods). Instead of assuming a diploid context like in the nuclear genome, the binomial parameter(s) here can range from 0 to 1 for different levels of heteroplasmy. The difference of the Bayesian Information Criterion scores of the fitted H_0_ or H_1_ models (ΔBIC=BIC(H_0_) - BIC(H_1_)) is then used to prioritise the candidate variants that are informative with respect to clonal discovery, with a higher ΔBIC for stronger support of the alternative model that the variant is clonally informative. Then, a cutoff on ΔBIC is determined automatically by the inflexion point (a knee point) in the cumulative distribution of log(ΔBIC) (Methods). With a highly discriminative set of mtDNA variants identified by MQuad, vireoSNP clusters single cells to clones based on their mtDNA mutation profiles.

**Fig. 1:**
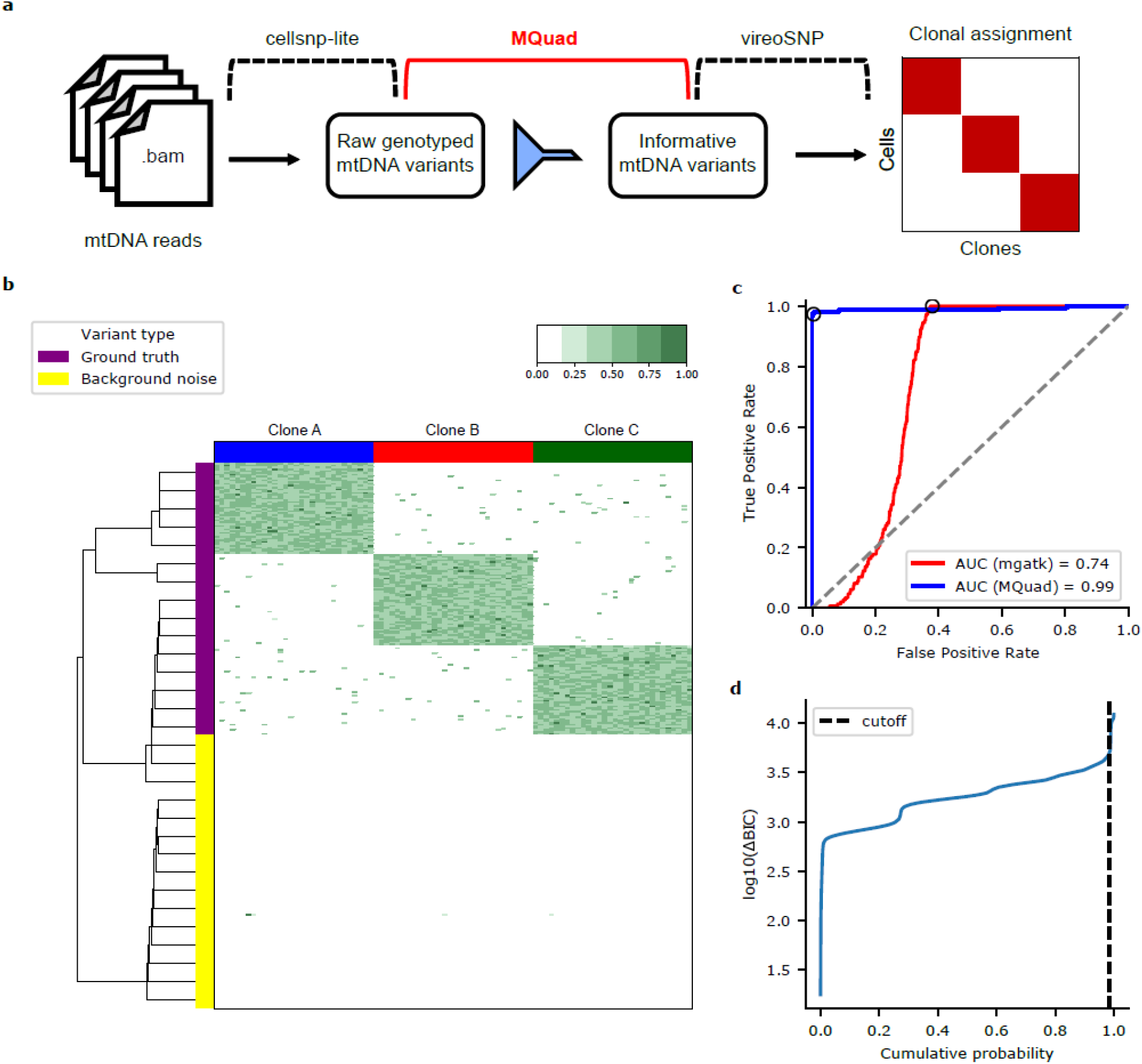
Overview of analysis pipeline and benchmark with simulated data. **a,** Schematic for tailored analysis suite recommended to use alongside MQuad. **b,** Simulation context for 90 cells in a hypothetical 3-clone structure. Each column is a cell, each row is a variant. Heatmap color represents mutated allele frequency. Only a small portion of background noise variants is shown for comparison with the actual clonal variants. **c,** Receiver operator characteristic curve showing the proportion of ground truth variants detected (true positive rate) as a function of proportion of background noise variants detected (false positive rate). Black circle represents the threshold used to classify informative variants. **d,** Cumulative distribution function of log(ΔBIC) with cutoff shown.

We first benchmarked MQuad with simulated data that mimicked high depth scRNA-seq data generated from the Smart-seq2 protocol. To simulate three clonal populations, we separated the dataset into three equal proportions and designated 50 clone-specific mtDNA variants for each population (Fig. 1b and Methods). On top of that, spontaneous mutations and sequencing errors were generated at a low frequency across all cells to simulate a noisy background (Methods).

MQuad achieved high accuracy in identifying ground truth clonal variants (area under receiver operator characteristic curve: AUC=0.99), outperforming the only other existing computational method mgatk (AUC=0.74) by a large extent (Fig. 1c). A high false positive rate was observed for mgatk (dot on blue ROC curve in Fig. 1c; FPR=0.38), consistent with the comments from the original publication^12^. We reason that the variance mean ratio (VMR) statistic used by mgatk is difficult to estimate and may suffer from high estimation variance, hence is not a robust predictor of informativeness in scRNA-seq due to a dominant presence of low allele frequency variants and sequencing errors. In contrast, our proposed metric, log(ΔBIC), showed distinct patterns between informative variants and noise, which made it more intuitive to place a cutoff at the knee point with the sharpest increase of log(ΔBIC) (Fig. 1d; dot on red ROC in Fig. 1c, FPR=0.005, TPR=0.973).

### Tumor cell populations are accurately inferred from informative mtDNA variants detected by MQuad

Next, we applied MQuad to a clear cell renal cell carcinoma (ccRCC) scRNA-seq dataset^20^ that contained a mixture of with three distinct source populations of cells from the same patient: patient-derived xenograft of the primary tumor (PDX pRCC); patient-derived xenograft of metastatic tumor (PDX mRCC); and metastatic tumor directly from the patient (Pt mRCC).

MQuad detected 133 informative mtDNA variants from the dataset. We observed heterogeneity in allele frequencies of these variants with some being highly clonal-specific, eg. 7207G>A was mostly specific to PDX pRCC alone and not found in the metastatic tumor populations (Fig. 2a). We also observed a sharp increase in the cumulative distribution of log(ΔBIC) (Fig. 2b) which was consistent with simulated data, further justifying the rationale behind determining the cutoff based on a knee point.

**Fig. 2:**
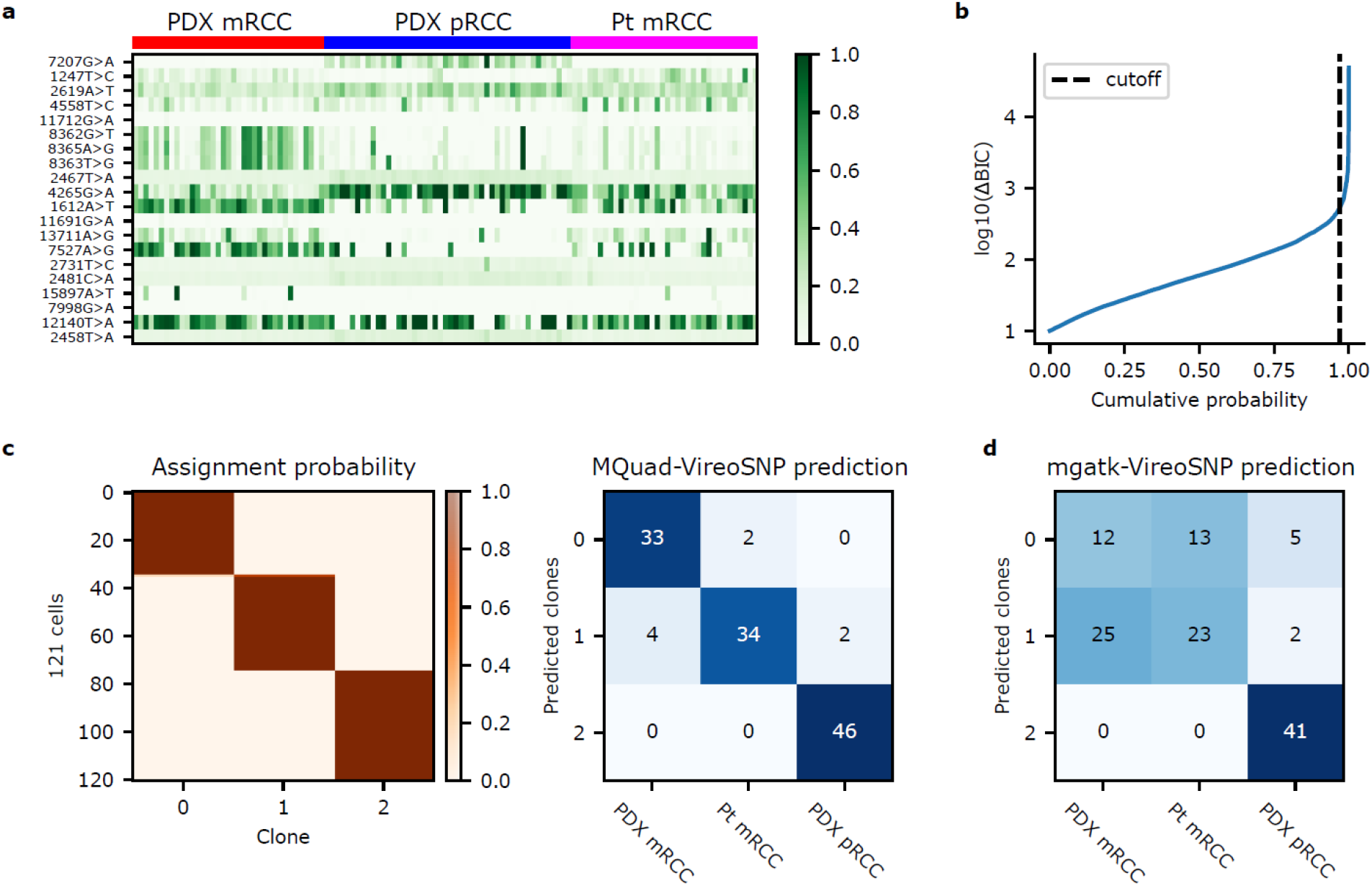
Distinct tumor populations inferred from heterogenous mtDNA variants. **a,** Allele frequency heatmap showing the top 20 informative mtDNA SNVs detected by MQuad ranked from highest to lowest ΔBIC. Each row is a variant, each column is a cell. **b,** Cumulative distribution function of log(ΔBIC) with cutoff shown. **c,** (Left) Clonal assignment with mtDNA variants detected with MQuad. Each row is a cell, each column is a clone, heatmap color indicates assignment probability. (Right) Confusion matrix between predicted mitochondrial clones and source labels. Numbers represent cells assigned. **d,** Confusion matrix between predicted clones based on mgatk variants and source labels.

To further validate the accuracy of MQuad, we used the detected variants as an input to vireoSNP for clonal assignment. We found that most cells were confidently assigned to their respective origins with high concordance (Fig. 2c). Our assignment achieved 91% concordance with the source labels, showing that genetic heterogeneity in the mitochondrial genome can indeed be leveraged for reconstruction of tumor clonal subpopulations.

We also assigned clones based on variants detected by mgatk. While mgatk detected more than double the number of variants (312 mtSNVs), the clonal assignment was actually less concordant (67%) with the source labels (Fig. 2d). The two metastatic tumor populations were less distinguishable from each other, possibly due to a large number of false positive mtSNVs that mgatk failed to filter out.

### MQuad identifies mtDNA-based clonal structure that complements clones inferred from nuclear SNVs

We further applied MQuad to a recent scRNA-seq dataset that characterized the somatic clones in healthy fibroblast cell lines^5^. The fibroblast cell line used, joxm, was from a white female aged 45-49. We observed even in cells that were not known to be tumorigenic, a substantial level of mitochondrial genetic heterogeneity could be detected. MQuad detected 59 SNVs with which we assigned the 78 cells into 3 clones (Fig. 3a). Not only did we observe distinct mutations in specific clones (e.g. 11196G>A, Fig. 3a and Supp. Fig. S3), we also identified random genetic drift events that lead to different heteroplasmy levels between clones (e.g. 2619A>T, Fig. 3a and Supp. Fig. S3).

**Fig. 3:**
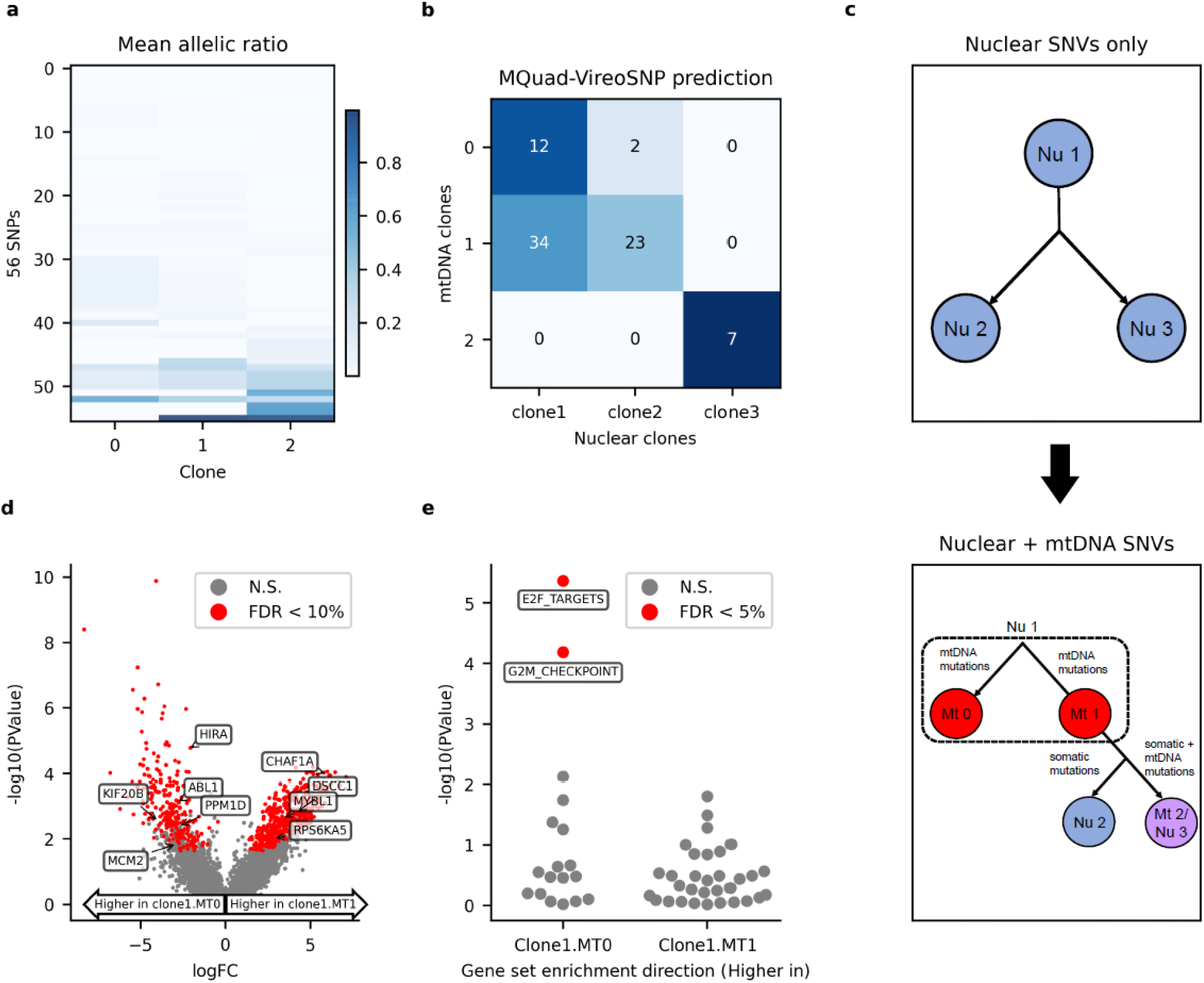
Combination of mtDNA and nuclear SNVs identifies finer clonal structure in healthy fibroblast. **a,** Mean allelic frequency of mtDNA variants in each clone. Each row is a mtDNA SNV, each column a clone. **b,** Confusion matrix between predicted mtDNA clones from MQuad and predicted nuclear clones from Cardelino. **c,** (Top) Clonal tree inferred from nuclear SNVs alone. (Bottom) Clonal tree inferred from both nuclear and mtDNA SNVs. (Nu: Nuclear clone; Mt: Mitochondrial clone) **d,** Volcano plot showing negative log10 *P*-values (edgeR two-sided quasi-likelihood *F-*test) against log2-fold changes (FC) for DE between cells assigned to Nu1 & Mt 0 and Nu1 & Mt 1. Significant DE genes (FDR < 0.1) highlighted in red. **e,** Enrichment of MsigDB Hallmark gene sets based on log2 FC between Nu1 & Mt 0 and Nu1 & Mt 1 (two-sided camera test). Gene set enrichment direction is Nu 1 & Mt 1 with respect to Nu 1 & Mt 0. Negative log10 *P*-values of gene set enrichments are shown with significant gene sets (FDR < 0.05) highlighted and labelled.

Comparing our clonal assignment to the clones inferred from nuclear mutations, mtDNA clones showed partial but consistent concordance with nuclear clones (Fig. 3b). In particular, nuclear clone 3 was completely identical to mitochondrial clone 2, while nuclear clone 1 and 2 were more ambiguous in terms of mtDNA. Using our MQuad-identified mtDNA variants, our pipeline discovered two novel subclones MT0 and MT1 within nuclear clone 1 that had distinct mtDNA mutation profiles (Fig. 3c). Next, we tested if MT0 and MT1 could indeed be biologically and clonally distinct cell populations. Differential expression (DE) analysis between cells from the two subclones of nuclear clone 1 (clone1.MT0 & clone1.MT1) identified 565 DE genes (two-sided edgeR QL *F*-test; FDR <0.1; Fig. 3d). A large number of highly expressed DE genes in clone1.MT0 were enriched for cell proliferation^21,22^ (gene sets E2F targets, G2M checkpoint; two-sided camera test; FDR < 0.1; Fig. 3e). The original study which published this dataset identified cells in clone 1 has a higher proliferation rate than clone 2. Using MQuad, our results further discovered that a specific subclone MT0 within clone 1 is likely the main contributor of the elevated proliferation rate.

### MQuad identifies subclonal structure in a gastric cell line with scDNA-seq

We further asked if mitochondrial mutations could be used in conjunction with copy number variations (CNVs) to infer clonal evolution in tumors. We first examined the applicability of MQuad on a barcode-based single-cell whole genome sequencing (scDNA-seq) dataset, e.g., 10x Genomics, as it had attracted a lot of attention recently owing to its accuracy of identifying CNVs and clonal structure despite its low coverage ^23,24^. By re-analysing a publicly available gastric cancer cell line MKN-45 (scDNA-seq in 10x Genomics) with both cellranger and our own B-allele frequency (BAF) clustering, we identified two distinct CNV clones (Fig. 4a, Supp. Fig. S3; Methods), which was in agreement with the original analysis ^23^. Both clones shared multiple CNVs, e.g. copy loss on chr4p, but clone 1 represented a small population of cells (7%) having a unique copy loss on chr4q.

**Fig. 4:**
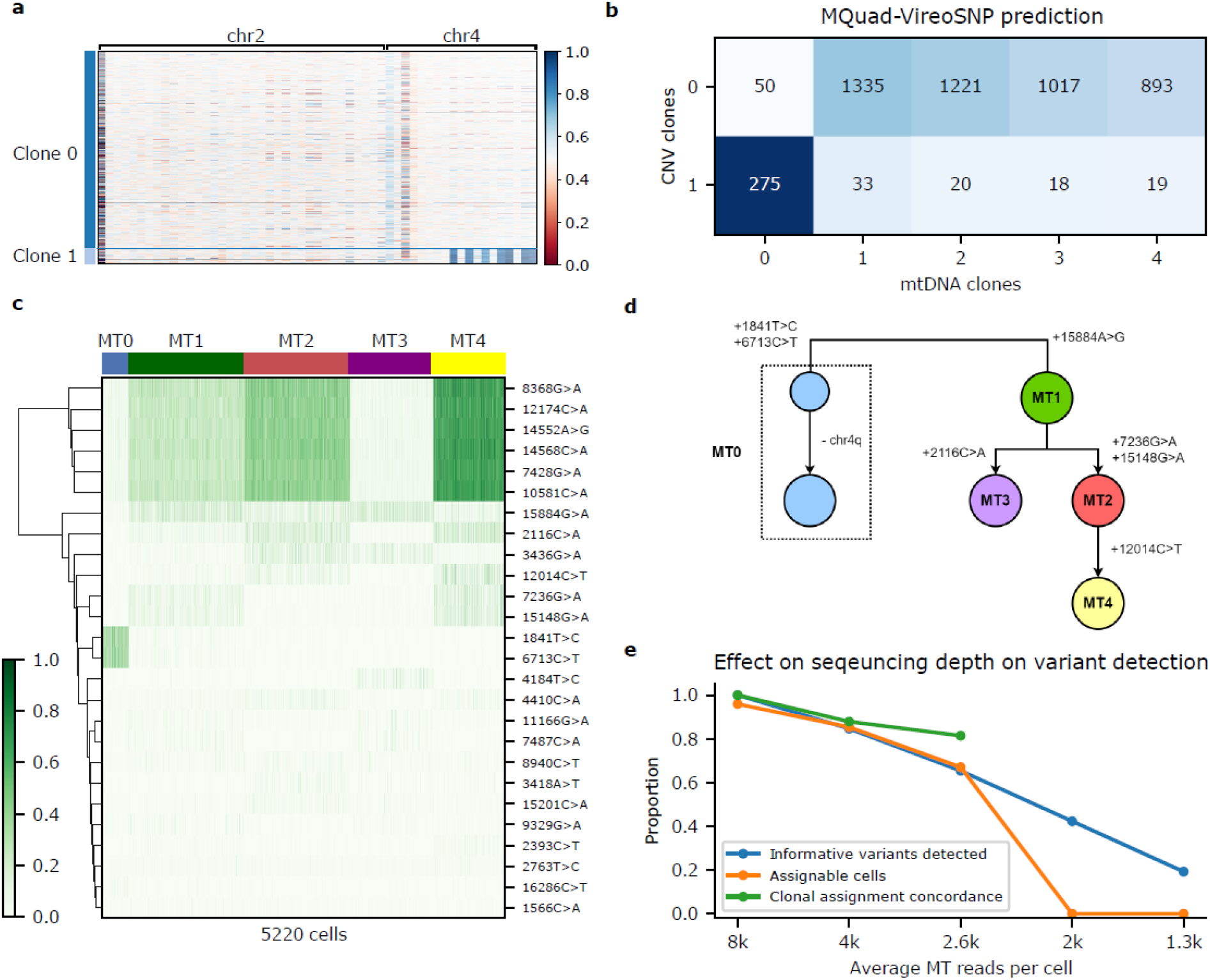
Mitochondrial mutations show concordance with CNV in scDNA-seq. **a,** CNV profile of gastric cancer cell line MKN-45. Shown is B-allele frequency across 5,220 cells (in row) on chr2 and chr4 on heterozygous SNPs aggregated in 5MB bins through phased in a three-step strategy (Methods). **b,** Confusion matrix between CNV clones and mitochondrial clones. **c,** Allele frequency heatmap on 26 clonally discriminative mtDNA variants detected by MQuad. **d,** A speculated lineage tree on CNV and mitochondrial clones considering the CN events and mtDNA allele frequency. **e,** Effects of sequencing depth on mtDNA variants identification and cell assignment to clones. For 2K and 1.3K reads per cell, clone structure cannot be detected by vireoSNP.

By applying MQuad to MKN-45 scDNA-seq, we detected 26 clone discriminative mitochondrial mutations (ΔBIC > 500) even with a relatively low sequencing depth and were able to infer 5 clones with vireoSNP (Fig. 4c). By comparing to CNV profiles, we observed a high degree of overlap between the CNV clone 1 and mitochondrial clone 0 (Fig. 4b, c). Based on the raw allele frequencies of mtDNA variants (Fig. 4c), we discovered the most distinctive clone of MT0 is marked by the presence of 1841T>C and 6713C>T, which were largely absent in all other clones (Fig. 4c). We also speculated a lineage tree that utilizes information from both CNVs and mtDNA SNVs (Fig. 4d).

Next, we downsampled the data into various depths to quantify the effect of sequencing depth on mtDNA variant calling and clonal assignment (Fig. 4e). Using the full dataset as a silver standard, we observed a significant drop in number of cells with assignable clones, number of informative mtDNA detected, and clonal assignment concordance when there were less than 2,600 mitochondrial reads per cell. Therefore we would recommend a minimum of 3,000 mitochondrial reads per cell for applying MQuad to perform adequately on scDNA-seq data, assuming that the reads were distributed evenly along the mitochondrial genome.

### MQuad can detect mtDNA variants near captured sites in UMI-based scRNA-seq

One of the most widely used platforms for scRNA-seq is droplet-enabled UMI-based scRNA-seq, such as those generated by the 10x Genomics platform. One of the main limitations of using such data for variant calling is that the reads are typically only enriched for the 3’ or 5’ end of a transcript, and hence resulting in reads that have highly non-uniform coverage. To test the performance of MQuad and the clonal assignment pipeline, we evaluated its application on three 3’-biased scRNA-seq datasets generated by the 10x Genomics platform from triple negative breast cancer samples (TNBC1, TNBC2 and TNBC5) ^10^. Unsurprisingly, even with a comparable number of mitochondrial reads, 10x scRNA-seq performed significantly worse than other sequencing protocols. Only a small number of mtDNA variants were found and the numbers of clones identified were small across all datasets (Supplementary Table S1). Coverage analysis showed this might be due to the highly uneven coverage of mitochondrial reads in this 3’-biased 10x Genomics scRNA-seq dataset (Fig. 5a). No or low number of cells can be confidently assigned to a clone in TNBC2 and TNBC5.

**Fig. 5:**
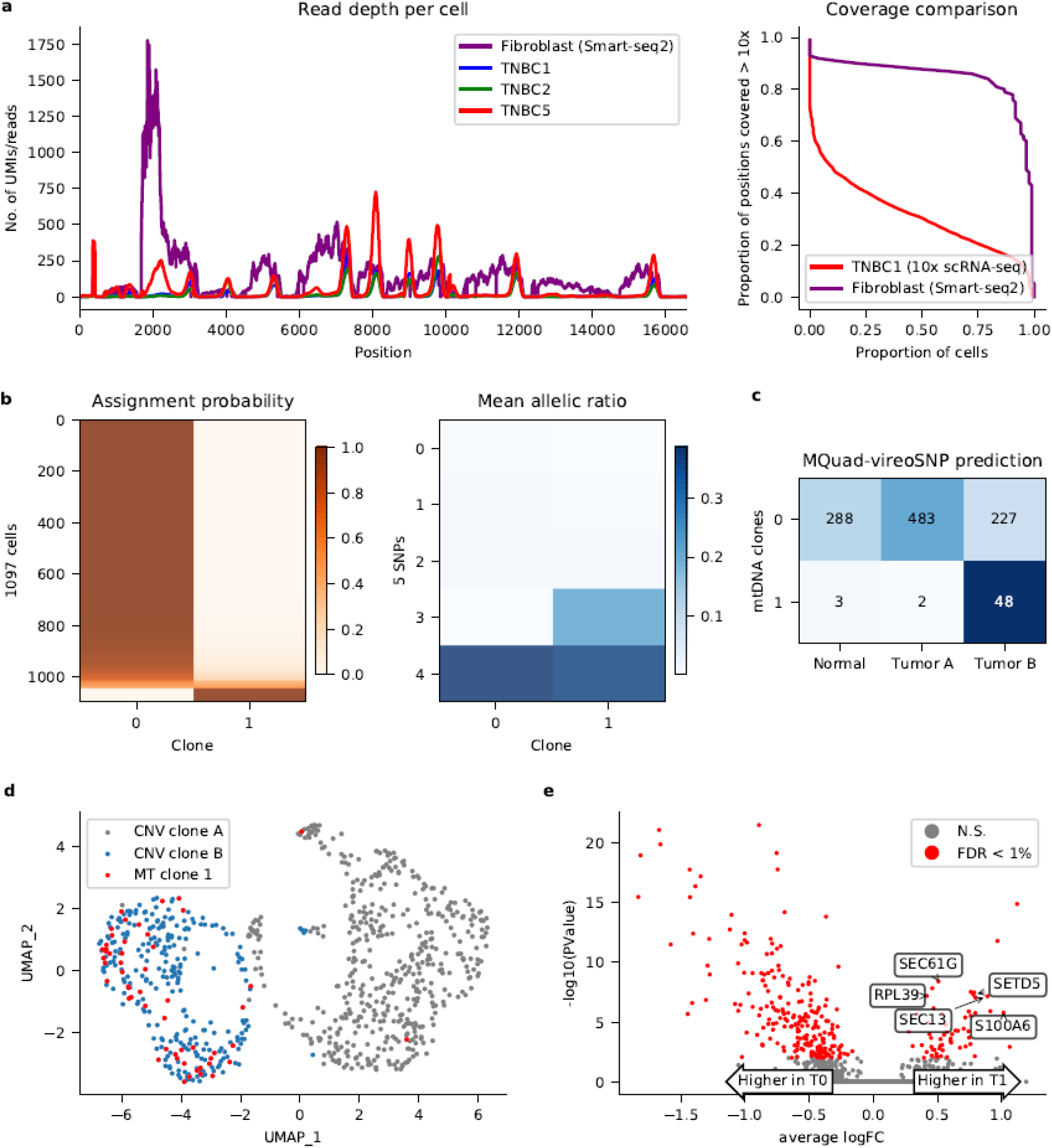
Analysis of mtDNA variants on 10x scRNA-seq datasets. **a,** Coverage comparison between 3 TNBC 10x scRNA-seq datasets and fibroblast Smart-seq2 dataset. UMI-based scRNA-seq shows uneven distribution of reads across the mitochondrial genome while Smart-seq2 is generally more well-covered. **b,** Clonal assignment and mean allelic ratio of 1097 cells from TNBC1. **c,** Confusion matrix between CNV subclones and confidently assigned mitochondrial clones. **d,** UMAP plot with tumor cells only. Mitochondrial clone 1 highly overlaps with CNV clone B. **e,** Volcano plot showing negative log10 *P*-values (Seurat FindMarkers with DESeq2 model) against average log fold changes (FC) for DE between T0 (Tumor & MT0) and T1 (Tumor & MT1). Significant DE genes (FDR < 0.01) highlighted in red. DE genes that are associated with breast cancer are labelled.

Nonetheless, MQuad detected five informative variants in the TNBC1 dataset and assigned 1051 cells (95.8%) with probability > 0.8 into two clones (Fig. 5b). In this case, the identified mtDNA variants were located close to the 3’ end of some genes. We found that the small mitochondrial clone 1 almost exclusively (50 out 53 cells) came from the tumor portion and showed a significant enrichment compared to mitochondrial clone 0 (p = 6 x 10^−5^, Fisher’s exact test; Fig. 5c). Moreover, mitochondrial clone 1 highly overlapped with CNV subclone B identified from the original paper^24^ (Fig. 5d), indicating the presence of a small subclone within a CNV clone. By examining the differentially expressed genes between the two mtDNA clones, we identified 57 up-regulated genes in subclone 1 (FDR < 1%, Seurat FindMarkers function with DESeq2 model; Fig. 5e), many of which are known to be associated with breast cancer, ie. SEC61G^25^ and RPL39^26^. High level of SETD5 expression is also related to poor prognosis in breast cancer^27^, providing additional evidence for the biological significance of mitochondrial clone 1.

Taken together, this analysis demonstrated the ability of MQuad to identify informative mtDNA variants to identify biologically meaningful cell subclonal structure, even in 3’ bias 10x Genomics data, suggesting that clonally-informative variants can be detected if they are close to the capture sites, which opens up a wide possibility to re-analyse a large volume of UMI-based scRNA-seq data in the public domain. Moreover, the technical issue of low read count and uneven coverage may be overcome by using emerging protocols for mitochondrial sequence enrichment^28^, further supporting the need for a highly effective computational method for detecting mitochondrial variants like MQuad and using such variants to detect the fine clonal structure in the cell population.

## Discussion

To date, mitochondrial sequences are often overlooked in single cell sequencing data analysis. We demonstrate here a novel analytical suite that can harness these mitochondrial sequences to discover clone-discriminative genetic variants using standard single cell sequencing data. Without requiring additional mitochondrial enrichment steps, our pipeline is applicable to most existing single-cell data with a sufficient sequencing depth and even coverage. This means that cell lineage information can be discovered with virtually no additional experimental cost. We make the MQuad source code and tool publicly available, and the program is designed to work seamlessly with other recently published tools, cellsnp-lite^18^ and vireo^19^. Our analysis shows that leveraging mtDNA mutations can decipher not only tumor clonal dynamics but also resolve clonal substructure that may not be detectable based on nuclear DNA alone.

The key strength of MQuad is that it adopts a model selection approach in evaluating the informativeness of each variant, which is more robust than considering the raw allele frequencies alone. Especially in deeply sequenced data with a lot of read counts (eg. Smart-seq2), this approach is more effective in reducing false positives compared to existing methods. With the emergence of massively parallel sequencing protocols that enriches mitochondrial reads, dealing with noise will be inevitable and MQuad serves as a flexible option to filter for useful mtDNA variants. Here we show that MQuad can be applied to sequencing data of various nature and we anticipate that it can adapt well to future sequencing technologies and datasets.

With the possibilities enabled by mtDNA lineage reconstruction, dealing with different types of genomic alterations occurring in the same cell remains an open challenge and there is a high demand for new computational methods for this purpose. In this study, we showed the correlation between mitochondrial and other types of genomic alterations such as CNVs and nuclear SNVs, but the correlation is mostly based on simple concordance and intuitive analysis. With MQuad, it is now possible to take mtDNA variations into account during retrospective lineage tracing, hence a more sophisticated model for the integration of CNV, mtDNA and nuclear SNVs will be highly beneficial for clonal analysis and lineage reconstruction.

To conclude, the methods presented here unlock the untapped potential of existing single-cell sequencing data and provide an effective approach in lineage reconstruction with mtDNA variants alone or together with nuclear SNVs and/or CNVs. As new sequencing technologies are evolving, MQuad opens up a new paradigm of analysis for a variety of single-cell sequencing datasets.

## Methods

### Variant detection pipeline

The tailored variant detection pipeline can be briefly divided into 3 major steps:

Firstly, raw reads from BAM files are piled up using **cellSNP-lite** (v1.2.0) ^18^. This generates a SNP-by-cell matrix in the form of a VCF file or sparse matrices of each cell’s AD and DP at each variant position. The output includes every SNP found in the mitochondrial genome, which contains a large amount of noise and uninformative variants. MQuad (v0.1.1) takes this output and selects high quality informative variants with a binomial mixture model (explained in the next section). Lastly, **vireoSNP** (v0.5.3) ^19^ uses variational inference to reconstruct clonal populations based on the selected SNPs from MQuad.

### MQuad model

MQuad assesses the heteroplasmy of mtDNA variants with a binomial mixture model. In this model, it is assumed that the number of reads (or UMIs) for alternate allele (AD) for a SNP follow a binomial distribution with total trials as the depth of both alleles (DP) and success rate depending on the presence or absence of a variant:

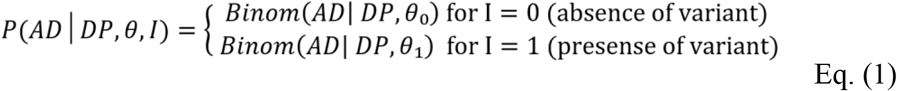

Assuming there are *M* cells in the sample, and the proportion of cells carrying a certain SNP is *π*, the likelihood can be estimated by:

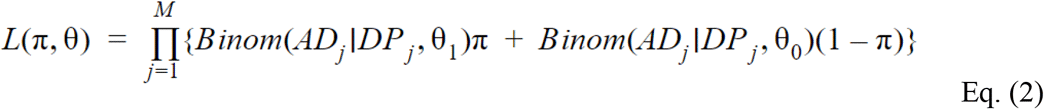

This likelihood of a 2-component binomial mixture model (M1) can be maximized using an expectation-maximization algorithm (pseudo code in Algorithm s1 in Supp. File) in order to get a maximum-likelihood estimation of π and θ. The same is done on a 1-component model (M0; ie. π = 0) via direct maximization. For each fitted model, the Bayesian Information Criterion (BIC) can be calculated with the obtained likelihood (*L*) and the penalty on the number of parameters (1 for M0 versus 3 for M1) by

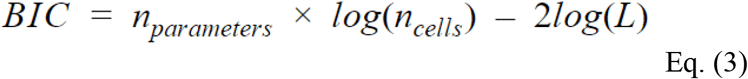

Consequently, the difference in Bayesian Information Criterion (ΔBIC) between models M1 and M0 can be calculated by ΔBIC = BIC (M0) - BIC(M1). The ΔBIC is further used as an indicator of clonal informativeness for each SNP with higher ΔBIC being more informative.

Finally, a ΔBIC cutoff for selection of SNPs is determined using the Kneedle algorithm^29^, from a Python package kneed (v0.7.0). Briefly, Kneedle defines the curvature of any continuous function *f* as a function of its first and second derivative:

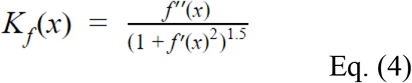

The algorithm aims to locate the ‘knee’ point by finding the point where *K_f_*(*x*) is maximum. This corresponds to our aim to find a point where the ΔBIC sharply increases to identify outlier SNPs which are most likely to be clonally discriminative.

### Data preprocessing

The scRNA-seq data (Kim and fibroblast datasets; both Smart-seq2) was preprocessed largely based on the guidelines from GATK: (https://gatk.broadinstitute.org/hc/en-us/articles/360035535912-Data-pre-processing-for-variant-discovery). Specifically, FASTQ files were first aligned to the reference genome (hg19 for human, mm10 for mouse) using STAR aligner^30^ with default parameters. Then, duplicates were marked and removed from the raw BAM files using MarkDuplicates, resulting in analysis-ready BAM files for variant detection.

For 10x data (MKN-45 scDNA-seq & TNBC scRNA-seq), analysis read BAM files are directly downloaded and used. It should be noted that they are aligned to hg38 reference, different from our processed data.

### Simulation

Mitochondria reads were synthesized in silico with NEAT-genReads v3.0 ^31^. Briefly, pair-end aligned reads were generated from a random mutation profile (average mutation rate = 0.05) for every simulated cell. A total of 90 cells with a 3-clone structure was simulated. The simulation was performed based on the standard hg19 reference genome, with a read length of 126bp, coverage of 300x per position per cell, and 1% sequencing error across all sites. We also designated 149 positions to insert clone-specific variants, with each clone having 49-50 variants. Since the mutation model of NEAT-genReads assumes a diploid context, all variants were simulated with 50% allele frequency.

### Analysis of copy number variations on scDNA-seq data

For MKN-45 scDNA-seq dataset, we first explored the copy number variation profile by CellRanger, a build-in software from 10x Genomics (Supplementary Fig. S3a). The CellRanger calling result shows a potentially interesting clonal structure with a unique small clone (node ID: 10050) carrying CN loss on chr4q, while a substantial fraction of cells and genomic regions may suffer from high error rate due to the high genomic variability in cancer cell line and the lack of allelic information. Therefore, we first confirmed that there is a genuine CN loss on chr4q in this group of cells by presenting its averaged B-allele frequency with comparison to the remaining cells (Supplementary Fig. S3b). In order to identify a cleaner clone assignment, we further used the read counts for both alleles on chr2 and chr4 to cluster cells into two clones with a binomial mixture model implemented in VireoSNP (Fig. 4a; Supplementary Fig. S3c). We noticed that with such BAF information, a two-cluster structure can be easily identified as suggested in the original study ^23^, which is also well concordant to the cellranger detected clusters (Supplementary Fig. S3d).

In order to visualise smoothed BAFs in single-cells, we used a three-level phasing strategy to aggregate multiple SNPs, as introduced in CHISEL^24^. First, allelic read counts were summed up for SNPs in a 50Kb block by reference based phasing with Sanger Imputation server. Second, we combined 100 blocks into a 5Mb bin by a shared allelic ratio. Third, the B alleles in near bins were flipped using a dynamic programming algorithm to achieve a minimal BAF discrepancy with neighbour bins. It should be noted that procedures of the aggregation across blocks (step 2) and allele flipping across bins (step 3) only contribute to the visualization in Fig. 4a, but do not affect the clustering of cells into CNV clones as they were directly based on the aggregated SNPs in a 50Kb window from reference based phasing (step 1).

### Gene expression analysis

Differential expression analysis on the fibroblast dataset was performed using the quasi-likelihood F-test function from edgeR^32^. To test for statistically significant differences in gene expression between clone1.MT0 and clone1.MT1, we fit a generalised linear model for single-cell gene expression with cellular detection rate, plate, and assigned mitochondrial clones as predictor variables.

Gene set enrichment analysis was performed using the camera function from limma^33,34^. Using 50 Hallmark gene sets from Molecular Signatures Database (MsigDB)^35^, we tested for their enrichment using the log2-fold-change from the previous edgeR model as input.

Both DE analysis and gene set enrichment steps were adjusted for multiple testing by FDR estimation using independent hypothesis weighting from IHW^36^. The independent covariate used was average gene expression.

Differential expression analysis on the TNBC1 dataset was performed using the FindMarkers function from Seurat^37^ using the DESeq2 model^38^.

### Data availability

All datasets used are all publicly available, through the NCBI Gene Expression Omnibus (GEO) portal or ArrayExpress database at EMBL-EBI, with accession numbers GSE73121 (Kim dataset), GSE148673 (TNBC datasets), E-MTAB-7167 (Fibroblast dataset).

MKN-45 scDNA-seq dataset is available from 10X Genomics website.

### Code availability

MQuad is an open-source Python package available at https://github.com/single-cell-genetics/MQuad. All the analysis notebooks for reproducing the results will be available on Zenedo once published.

## Acknowledgements

We thank Xinajie Huang for optimising cellSNP-lite on genotyping mitoDNA variants and preprocessing MKN-45 data, and Yin Hei Lam for reproducing CopyKat results on TNBC1 data.

