## Supplementary material for "MQuad enables clonal substructure discovery using single cell mitochondrial variants"

### Supplementary figures

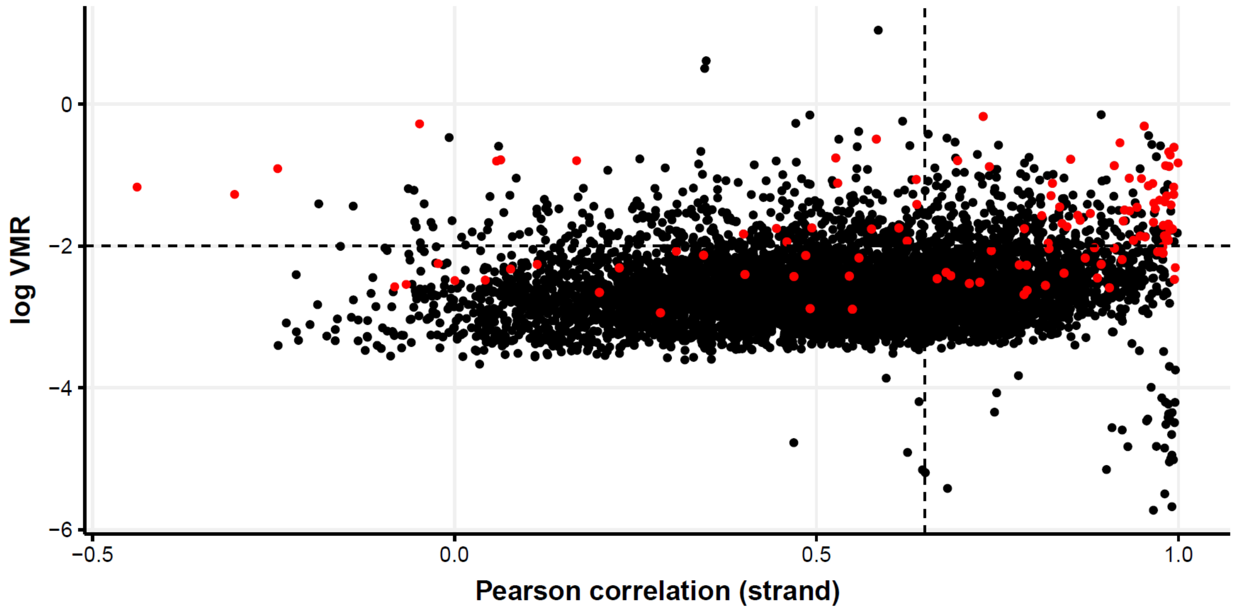


**Supplementary Fig. S1: Visualization of mgatk output for Kim dataset.**
Each dot represents a mtDNA variant, MQuad variants highlighted with red. Dotted lines are default thresholds for informativeness.

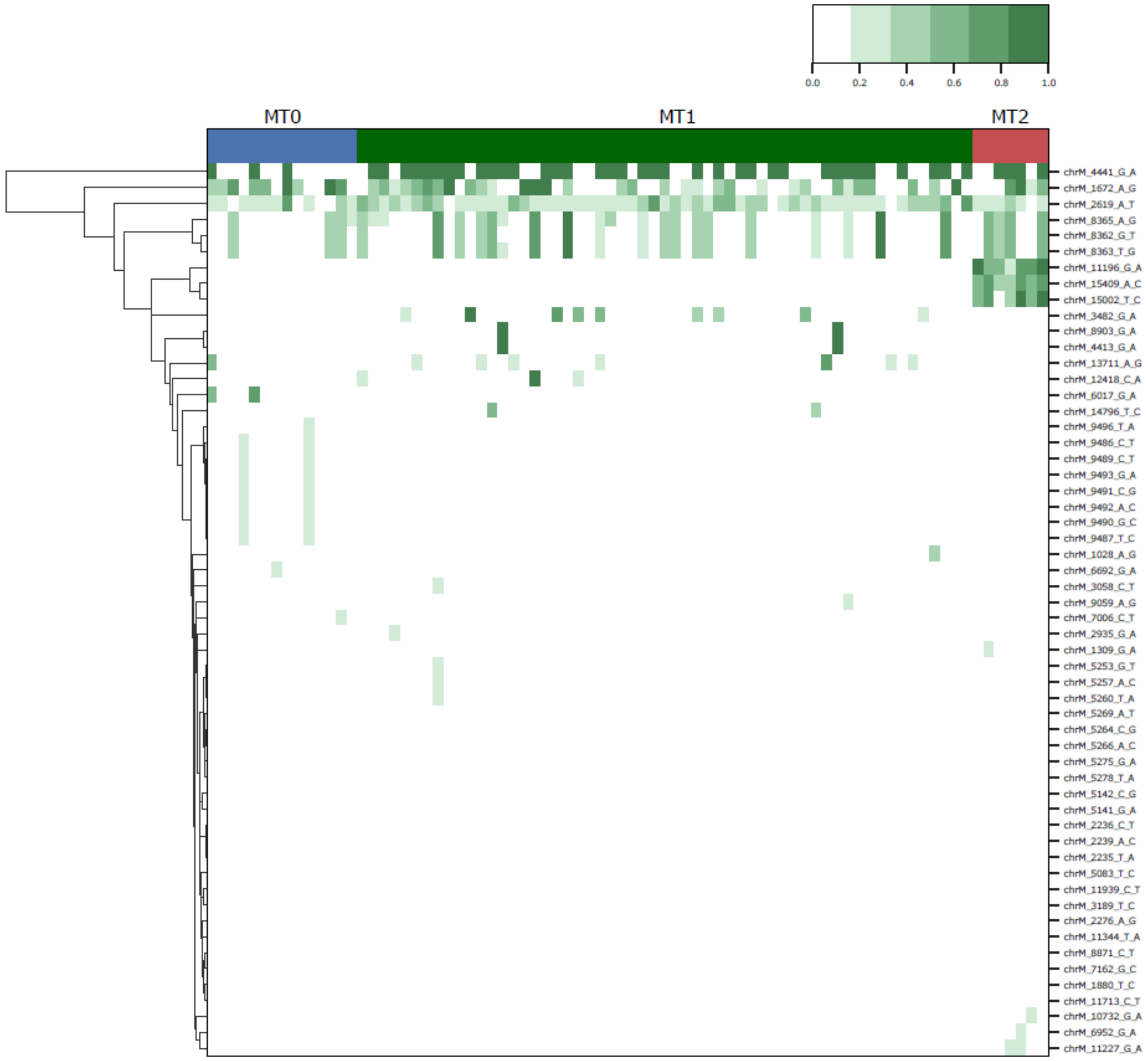


**Supplementary Fig. S2: Allele frequency heatmap of fibroblast dataset.**

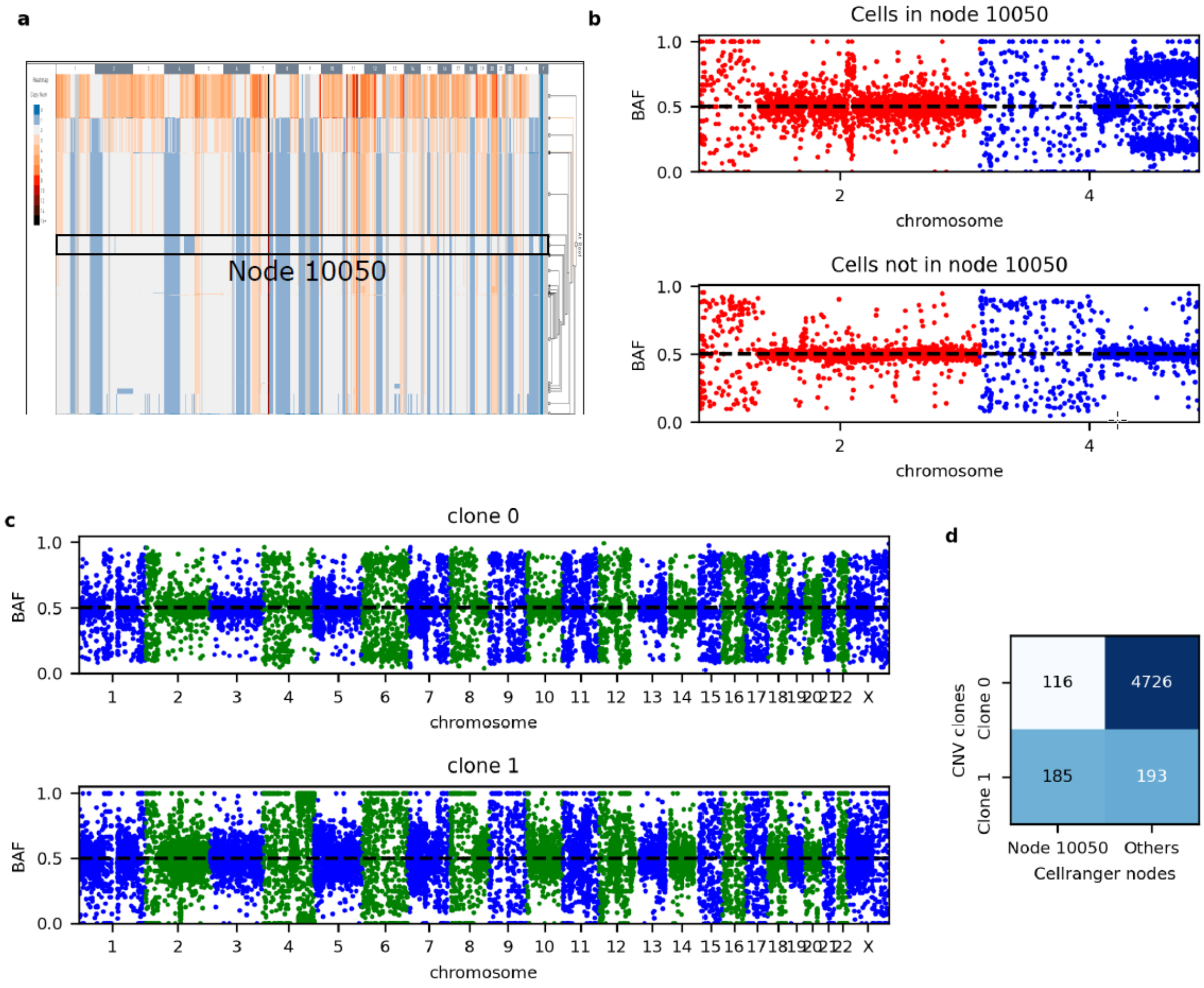


**Supplementary Fig. S3: Averaged B-allele frequency for CNV clones identified by VireoSNP binomial mixture model.**

**a,** Cellranger CNV profile of MKN-45 cell line. Node 10050 highlighted.

**b,** Averaged B-allele frequency (BAF) on chr2 and 4 for cells in node 10050 (top) and cells not in the node (bottom).

**c,** Averaged BAF on all chromosomes for CNV clone 0 (top) and clone 1 (bottom) that are identified with BAF on chr2 and chr4 (see Methods).

**d,** Confusion matrix between assigned CNV clones and original Cellranger nodes.

| Dataset | No. of cells | Average total reads per cell | Average MT reads per cell | No. of informative MT variants | Proportion of assignable cells | No. of clones |
|---|---|---|---|---|---|---|
| TNBC1 | 1,097 | 207,222 | 9,843 | 5 | 96% | 2 |
| TNBC2 | 1,034 | 139,018 | 6,817 | 3 | 0% | NA |
| TNBC5 | 3,225 | 99,940 | 14,953 | 16 | 5% | 2 |

**Supplementary Table S1: Summary of scRNA-seq datasets used in Fig. 5.**

**Algorithm 1:** Expectation Maximization

**Input:** AD, DP, K=2

**Output:** $\pi_k, \theta_k, (k = 0, .., K - 1)$

**for** *each k* **do**

Randomly Initialize $\pi_k, \theta_k$,

$s.t. \sum_k \pi_k = 1;\ 0 \leq \theta_k \leq 1$

**end**

**while** *Not Convergence* **do**

*# E-step*:

**for** *each k* **do**

**for** *j=1:M* **do**

$$\bar{\gamma}_k^{(j)} := \frac{\theta_k^{AD_j}(1-\theta_k)^{(DP_j - AD_j)}.\pi_k}{\sum_k \theta_k^{AD_j}(1-\theta_k)^{(DP_j - AD_j)}.\pi_k}$$

**end**

**end**

*# M-step*:

**for** *each k* **do**

$$\pi_k = \frac{\sum_{j=1}^{M} \bar{\gamma}_k^{(j)}}{N};\ \theta_k = \frac{\sum_{j=1}^{M} AD_j \bar{\gamma}_k^{(j)}}{\sum_{j=1}^{M} DP_j \bar{\gamma}_k^{(j)}}$$

**end**

**end**

**Supplementary Algorithm S1: Expectation Maximization**
